# Exploitation of the cooperative behaviours of anti-CRISPR phages

**DOI:** 10.1101/574418

**Authors:** Anne Chevallereau, Sean Meaden, Olivier Fradet, Mariann Landsberger, Alice Maestri, Ambarish Biswas, Sylvain Gandon, Stineke van Houte, Edze R. Westra

**Affiliations:** ESI, Biosciences, University of Exeter, Cornwall Campus, Penryn TR10 9EZ, UK.; Department of Microbiology and Immunology, University of Otago, PO Box 56, Dunedin 9054, New Zealand.; CEFE UMR 5175, CNRS Université de Montpellier Université Paul-Valéry Montpellier EPHE, 34293 Montpellier Cedex 5, France.

**Author notes:** MRC Laboratory for Molecular Cell Biology, University College London, London, UK.

## Abstract

Many bacteria encode CRISPR-Cas (<u>C</u>lustered <u>R</u>egularly <u>I</u>nterspaced <u>S</u>hort <u>P</u>alindromic <u>R</u>epeats; <u>C</u>RISPR-<u>as</u>sociated) adaptive immune systems to protect themselves against their viruses (phages)^1^. To overcome resistance, phages have evolved anti-CRISPR proteins (Acr), which naturally vary in their potency to suppress the host immune system and avoid phage extinction^2,3,4,5^. However, these Acr-phages need to cooperate in order to overcome CRISPR-based resistance^4,5^: while many initial infections by Acr-phages are unsuccessful, they nonetheless lead to the production of Acr proteins, which generate immunosuppressed cells that can be successfully exploited by other Acr-phages in the population^4,5^. Here we test the prediction that phages lacking *acr* genes (Acr-negative phages) may exploit this cooperative behaviour^6^. We demonstrate that Acr-negative phages can indeed benefit from the presence of Acr-positive phages during pairwise competitions, but the extent of this exploitation depends on the potency of the Acr protein. Specifically, “strong” Acr proteins are more exploitable and benefit both phage types, whereas “weak” Acr proteins predominantly benefit Acr-positive phages only and therefore provide a greater fitness advantage during competition with Acr-negative phages. This work further helps to explain what defines the strength of an Acr protein, how selection acts on different Acr types in a phage community context, and how this can shape the dynamics of phage populations in natural communities.

---

To explore the hypothesis that cooperative behaviours of Acr-phages may be exploited by Acr-negative phages in the environment, we first performed a theoretical analysis (see methods for a full description of the model). This model assumes that bacteria are initially sensitive (W) but can evolve CRISPR-based resistance upon lytic phage infection (Fig. 1A). Infection of a CRISPR-resistant bacterium (R) by a lytic phage encoding an Acr protein (hereafter referred to as “Acr-positive phage”) can lead either to cell lysis (V) with probability *ϕ*, or result in a failed infection that leaves the host in an immunosuppressed state (S), which reverts back to the resistant state at rate *γ*. Hence, *ϕ* and *γ*^−1^ (the average duration of immunosuppressed state) are both measures of the “strength” of the Acr, but with distinct social implications for other phages in the population: *ϕ* is best described as a selfish trait (i.e. the greater *ϕ*, the higher the probability that the initial phage will lyse the host and replicate), whereas *γ*^−1^ could be viewed as an altruistic trait (i.e. the greater *γ*^−1^, the greater the opportunity for other phages to exploit immunosuppressed cells, but with no direct benefit to the initial Acr-phage that caused the failed infection).

**Fig. 1.**
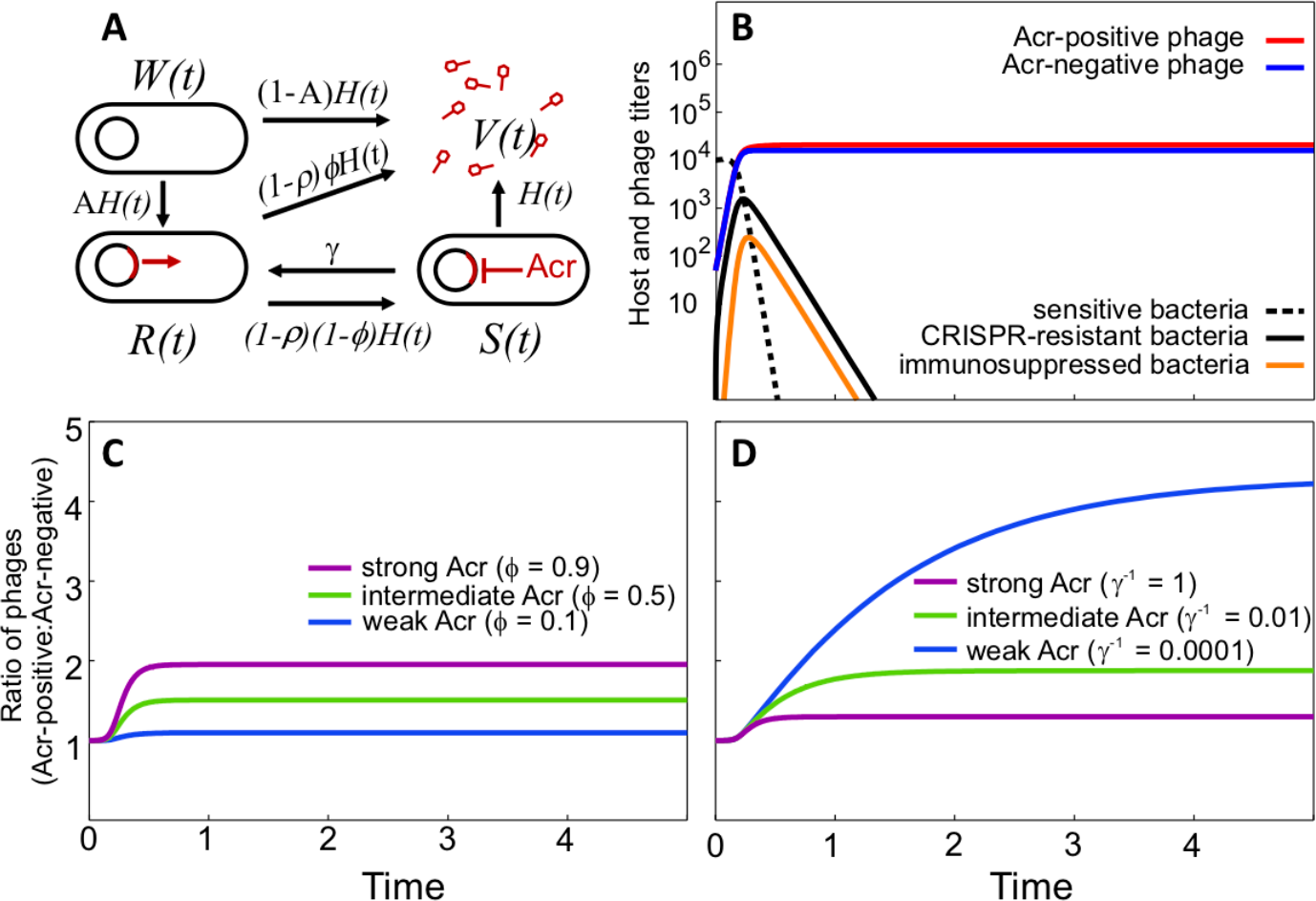
Altruistic and selfish effects of strong Acr proteins on Acr-negative phages. (**A**) Infection model of the Acr-phage (see details of the model in Supplementary Text). The parameter *H(t)* = a*V(t)* refers to the rate at which bacteria are infected by free phage particles, A is the probability that bacteria acquire CRISPR-based resistance. (**B**) Bacteria and phage population dynamics upon infection of initially sensitive hosts (dashed line) with an equal mix of 100 Acr-negative (blue line) and 100 Acr-positive phages (red line, *ϕ* = 0.3, *γ*^−1^=1). Initially sensitive bacteria evolve CRISPR-resistance (solid black line), which in turn can become immunosuppressed upon infection by Acr-positive phages (orange line). (**C**) Effect of *ϕ* and (**D**) *γ*^−1^ on the ratio of Acr-positive and Acr-negative phages following infection of sensitive bacteria. Other parameter values: a = 0.001, A = 0.2, B = 5, *ρ* = 0.5

As expected, this model predicts that infection of initially sensitive bacteria with lytic Acr-negative phages leads to rapid phage extinction due to the evolution of CRISPR-based resistance, whereas lytic Acr-positive phages avoid extinction regardless of the strength of their Acr (Fig. S1A-G). However, when sensitive bacteria are infected with an equal mix of the Acr-positive and Acr-negative phages, both phages avoid extinction (Fig. 1B). Interestingly, in this context, phages encoding stronger Acr proteins do not necessarily have a greater fitness advantage: the fitness of Acr-positive phages increases if stronger Acr proteins are more effective in directly lysing CRISPR-resistant hosts (i.e. when increased strength of Acr proteins is driven by higher values of *ϕ*) (Fig. 1C, S1H-J and S2), but decreases if stronger Acr proteins are more effective in maintaining CRISPR-resistant cells in an immunosuppressed state (i.e. when increased strength of Acr proteins is driven by higher values of *γ*^−1^) (Fig. 1D, S1K-M and S2). These opposing impacts of *ϕ* and *γ*^−1^ on phage relative fitness during pairwise competition can therefore help to elucidate whether the variations in Acr strengths originate from differences in their selfish ability to directly by-pass CRISPR-Cas immune systems, or from differences in their potential for altruistic cooperation.

To test these theoretical predictions, we set up an experimental system of mixed infections with Acr-positive and Acr-negative phages. To this end, we used *Pseudomonas aeruginosa* wildtype (WT) strain PA14 and the non-lysogenic phage DMS3*vir* which encodes an Acr protein (AcrIE3) that does not block the type I-F CRISPR-Cas system carried by the host (hereafter referred to as the Acr-negative phage). The WT strain naturally possesses spacers with only a partial match to the DMS3*vir* genome, and hence is initially phage-sensitive but primed to acquire targeting spacers, which leads to the rapid evolution of CRISPR-resistance^3,7,8^. As Acr-positive phages, we used isogenic versions of DMS3*vir* (namely DMS3*vir*-*acrIF1* and DMS3*vir*-*acrIF4*, described in Ref. 5), which encode Acr proteins specifically blocking the type I-F CRISPR-Cas system, instead of the native AcrIE3-coding sequence. Of these two phages, DMS3*vir*-*acrIF1* has a greater ability to amplify on pre-immunized bacteria (i.e. AcrIF1 is a “strong” Acr, and AcrIF4 is a “weak” Acr), a difference that can in principle be explained either by higher *γ*^−1^or higher *ϕ* (see ref. 5).

In accordance with our model predictions (Fig. S1B-G), both types of Acr-positive phages have similar population dynamics upon infection of initially sensitive WT bacteria (Fig. S3A). We verified the model assumption that WT bacteria could evolve CRISPR-based resistance against Acr-positive phages by performing deep sequencing analysis of host CRISPR loci at 3 days post infection (dpi). We found that spacers were indeed acquired following infection with DMS3*vir*-*acrIF4* or DMS3*vir*-*acrIF1*, at lower frequencies (Fig. S3B) but with similar patterns (i.e. number and location of acquired spacers) to those observed during evolution of CRISPR-resistance against Acr-negative phages (Fig. S3C-D). While this capacity to evolve CRISPR-resistance normally provides significant fitness benefits to bacteria when exposed to Acr-negative phages, it was no longer the case in the presence of Acr-positive phages, as measured in a 3-day competition experiment between the WT strain and an isogenic host lacking a functional CRISPR-Cas system (hereafter referred to as ‘CRISPR-KO’) (Fig. S2E).

Having confirmed that the phages behave consistently with our model predictions (Fig. S1A-G), we then used this empirical system to study the impact of strong versus weak Acr proteins during competition between Acr-positive and Acr-negative phages. We first verified that Acr-negative and Acr-positive phages performed equally well on the CRISPR-KO host, to ensure that any effect we may observe during competition solely depends on the strength of the Acr and not on any potential fitness cost inherent to a specific *acr* gene (Fig. 2A). Next, we co-infected the WT strain with a 50:50 mix of the Acr-negative and one or the other Acr-positive phage. After a 3-day competition (to allow time for evolution of CRISPR-resistance), we found that phages encoding the strong AcrIF1 had only a relatively small fitness advantage over Acr-negative phages (Fig. 2B, mean relative fitness *w*=2.0) whereas phages encoding the weak AcrIF4 more markedly outcompeted Acr-negative phages (Fig. 2B, *w*=9.8). This effect is consistent with the predictions associated with a high value of *γ*^−1^ in the model (Fig. 1D), which also predicts that the bacterial sub-population evolving CRISPR-resistance upon a mixed infection with Acr-positive and Acr-negative phages would be depleted more rapidly if the Acr is strong (Fig. S1K-M, black lines). Consistent with this prediction, we found that the proportion of bacteria with evolved CRISPR-resistance at 3 dpi was lowest when exposed to a mix of Acr-negative and strong AcrIF1-phages (Fig. 2C). To further validate the model prediction, we competed CRISPR-resistant bacteria (carrying 1 targeting spacer, hereafter referred to as ‘BIM-1sp.’) against surface-resistant bacteria in the presence of one or the other type of Acr-positive phages. Surface-resistant bacteria have evolved phage resistance through the loss of phage receptor^7^, which provides a protection equally effective against Acr-negative and Acr-positive phages. This experiment confirmed that the selection against CRISPR-resistant bacteria was indeed stronger in the presence of strong Acr-phages compared to weak Acr-phages (Fig. 2D).

**Fig. 2.**
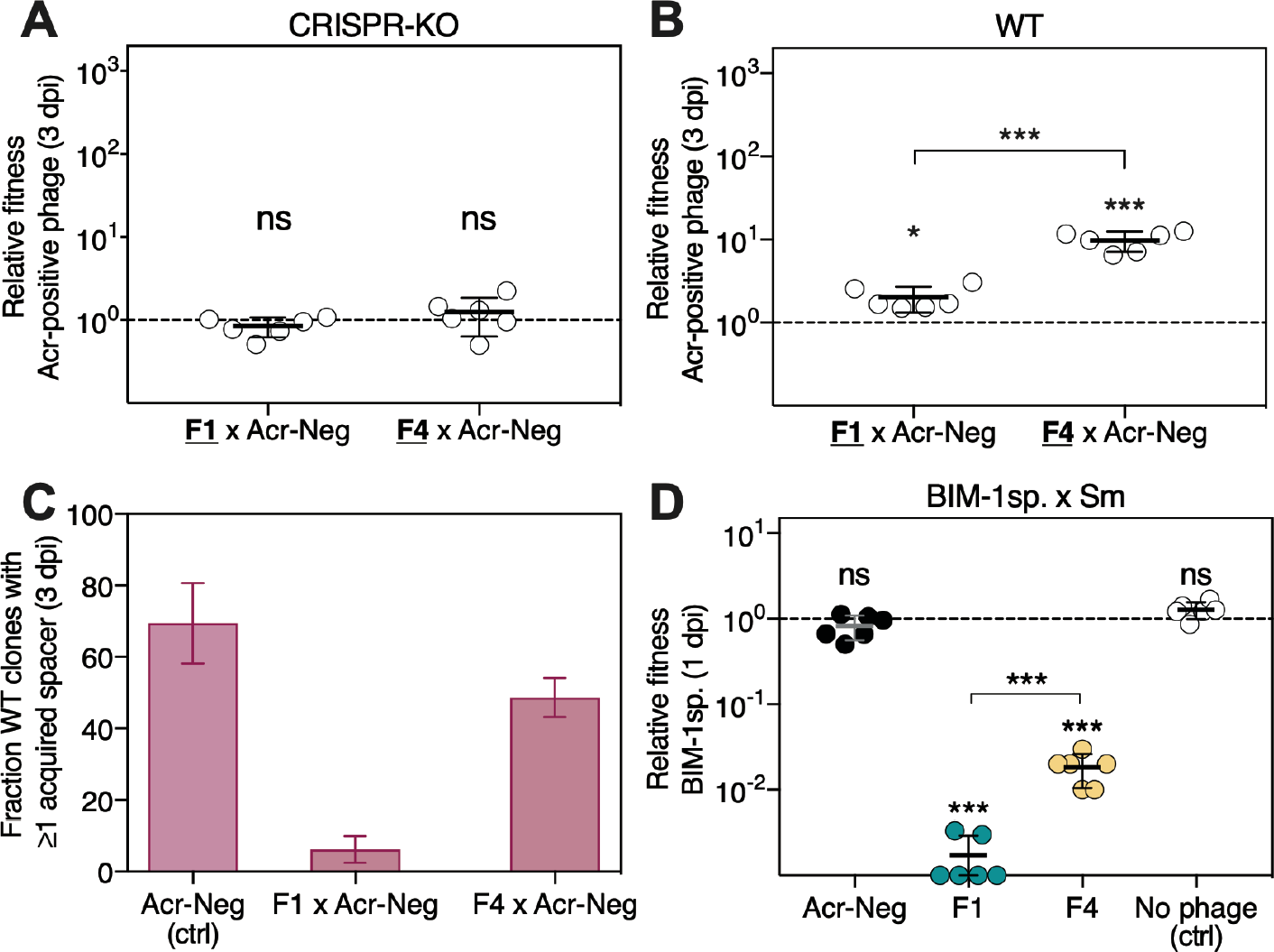
Low competitive advantage of strong Acr-phages over Acr-negative phages is linked to their stronger ability to deplete CRISPR-resistant clones from the host population. (**A**) Relative fitness of Acr-positive phages (as indicated by the underscored and bold font on the x-axis) upon 3-day competition on the CRISPR-KO strain (‘ns’ indicates no significant difference from 1: ‘F1 × Acr-neg’: p=0.15, t_5_=1.71; ‘F4 × Acr-neg’: p=0.3, t_5_=1.05) or (**B**) on the WT strain (stars indicate significant differences from 1 (‘F1 × Acr-neg’: p=0.01, t_5_=3.78; ‘F4 × Acr-neg’: p=0.0003, t_5_=8.63) or in-between values (p<0.0001, t_10_=7.40)). (**C**) Evolution of CRISPR-based resistance in the WT host population at 3 dpi upon individual or mixed infections (n=144; 24 clones tested in each of 6 independent infections). (**D**) Relative fitness of bacteria with CRISPR-resistance (BIM-1sp.: 1 targeting spacer) and surface-resistance (Sm) in the presence of indicated phages (MOI ~ 25). Significant differences from 1 (‘F1’: p<0.0001, t_5_=2189, ‘F4’: p<0.0001, t_5_=319) or in-between values (‘F1’ versus ‘F4’: p=0.0003, t_10_=5.35) as well as non-significant differences (‘Acr-Neg’: p=0.16, t_5_=1.67, ‘No phage’: p=0.05, t_5_=2.52) are indicated. All panels show values for individual replicates and/or means and 95% confidence intervals (n=6). ‘F1’=DMS3*vir*-*acrIF1,* ‘F4’*=*DMS3*vir*-*acrIF4*, ‘Acr-Neg’=DMS3*vir*.

Apart from this ecological effect of CRISPR-resistant host depletion, the smaller fitness advantage of strong AcrIF1-phages over Acr-negative phage also suggests that AcrIF1 causes a greater accumulation of immunosuppressed cells than AcrIF4, which can be exploited by Acr-negative phages (Fig. S1K-M and FigS2D-F, orange lines). To test this hypothesis, we competed Acr-positive and Acr-negative phages on pre-immunized BIM-1sp. strain with an initial 50:50 ratio and a multiplicity of infection (MOI) sufficiently high (~50) to enable Acr-positive phages to amplify on the CRISPR-resistant host. As expected, Acr-positive phages had a much higher fitness than Acr-negative phages, but this fitness advantage was approximately 10-fold greater for phages encoding the weak AcrIF4 (Fig. 3A, also see Fig. S2H). While densities of Acr-negative phages normally decreased by ~10^4^-fold at 1 dpi when grown individually on the BIM-1sp. strain, their densities decreased by only ~5-fold in the presence of the weak AcrIF4-phages and even increased by 3-fold in the presence of the strong AcrIF1-phage (Fig. 3B), hence providing direct evidence that DMS3*vir-acrIF1* generates higher amounts of immunosuppressed bacteria that can be exploited by Acr-negative phages. This ‘AcrIF1-assisted exploitation’ of CRISPR-resistant hosts by Acr-negative phages was no longer observed when the probability that immunosuppressed cells get re-infected before reverting to their resistant state was too low (i.e. at low MOI), further supporting the idea that Acr-negative phages take advantage of the mechanism of cooperative infections used by Acr-positive phages (Fig. 3C).

**Fig. 3.**
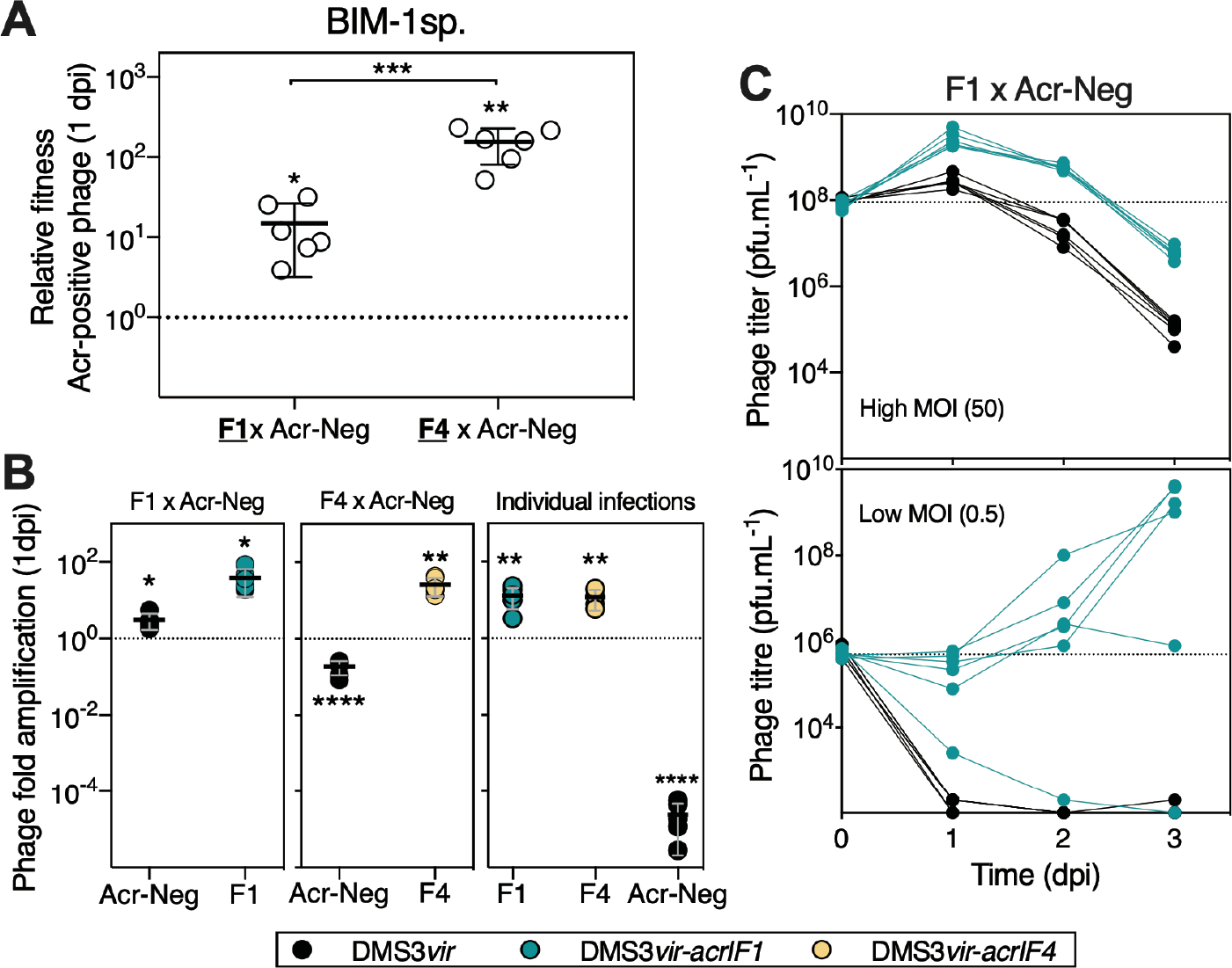
High immunosuppression mediated by strong Acr-phages allows exploitation of CRISPR-resistant bacteria by Acr-negative phages. (**A**) Relative fitness of Acr-positive phages (as indicated by the underscored, and bold font on the x-axis) upon 1-day competition on the CRISPR-resistant host (BIM-1sp). Significant differences from 1 (‘F1 × Acr-neg’: p=0.03, t_5_=3.05; ‘F4 × Acr-neg’: p=0.003, t_5_=5.39) or in-between values (p=0.0007, t_10_=4.84) are indicated. (**B**) Amplification of phages at 1 dpi upon either mixed or individual infections (as indicated on the top of the graphs) of the CRISPR-resistant BIM-1sp. host. Infections were performed with an initial MOI of 25. Stars indicate significant differences from 1 (‘F1 × Acr-neg’: (F1) p=0.01, t_5_=3.7 (Acr-Neg) p=0.01, t_5_=3.8; ‘F4 × Acr-neg’: (F4) p=0.0044, t_5_=4.9 (Acr-Neg) p<0.0001, t_5_=27.1; ‘Individual infections’: (F1) p=0.008, t_5_=4.2 (F4) p=0.008, t_5_=4.3 (Acr-Neg) p<000.1, t_5_=1.1×10^5^). (**C**) Phage population dynamics of F1 and Acr-Neg during mixed infection at initially high (top panel) or low (bottom panel) MOI. In all panels, values for individual replicates and/or means and 95% confidence intervals are indicated (n=6). ‘F1’=DMS3*vir*-*acrIF1,* ‘F4’*=*DMS3*vir*-*acrIF4*, ‘Acr-neg’=DMS3*vir*.

Altogether, these results revealed that phages with *acr* genes can benefit co-existing Acr-negative phages. Naturally, the population dynamics during mixed infections of Acr-positive and Acr-negative phages will inevitably depend on the level of cross-reactivity of the CRISPR-Cas immune system against the competing phages. A recent study analysing the CRISPR arrays from >700 *P. aeruginosa* genomes has shown that a given spacer usually provides such cross-reactivity as it typically matches several viruses (e.g. 2.75 viruses on average), which generally are genetically similar and co-occur in the same ecological niche (hence are likely to compete for the same hosts), but some spacers have been found to provide cross-reactive immunity against distantly related viruses^9^. The model and data presented here also show that during competition with Acr-negative phages, the strength of the Acr protein is a key determinant of the fitness benefits it provides. Notably, the greater benefits provided by phages encoding strong Acr proteins result from two effects. First, they deplete the pool of CRISPR-resistant hosts more rapidly than weak Acr-phages, as a result of their “selfish” interest to replicate on CRISPR-resistant hosts, which is somewhat analogous to the way antibiotic resistant bacteria can support growth of sensitive species by detoxifying the environment^10–12^. Second, the strong AcrIF1 induces longer periods of immunosuppression than the weak AcrIF4, which is consistent with data showing that AcrIF1 binds the Csy surveillance complex with higher affinity and slower off-rates than AcrIF4^4^. This “altruistic” production of immunosuppressed cells can be viewed as a “public good” that can be exploited by Acr-negative phages, similarly to iron scavenging molecules, communication signals and virulence factors, which are produced by few individuals but can benefit the whole population^13–15^. However, the evolutionary stability of such altruistic cooperation can be undermined by the invasion of cheats that do not contribute to public good production but still share the benefits^16^. Cooperation can be stabilized if the producers gain a greater proportion of the benefits (e.g. when diffusion of public goods is limited^17,18^), or if mutation of the cooperative gene carries large pleiotropic costs^19^. This latter mechanism could drive the evolution of Acr-cooperation if the abilities to directly lyse CRISPR-resistant hosts and to induce lasting immunosuppression are linked and therefore cannot evolve independently^20^. Alternatively, if these traits are independent, the observed cooperative behaviour is more likely driven by kin selection^16^, which is favoured by the spatially structured nature, and resulting high relatedness, of many bacterial and phage populations^21,22^. Future tests of these ideas are needed to fully understand how pleiotropy and kin selection drive the evolution of *acr* genes with different strengths and the observed variation in their cooperative traits, which has key implications for the ability of Acr-negative phages to amplify in the face of bacteria with CRISPR-Cas immune systems.

## Methods

### Mathematical modelling

We extended our previously described epidemiological model^5^, to model the population dynamics of bacteriophages that encode anti-CRISPR proteins (Acr) in an initially sensitive host population that can evolve CRISPR resistance. All simulations were performed in the software Mathematica version 11.2.

Bacteria may either be sensitive (the density of these bacteria is denoted *W*(*t*)), or may have evolved CRISPR-resistance after the incorporation of a spacer targeting the phage (the density of these bacteria is noted *R*(*t*)), or may be in an immunosuppressed state (the density of these bacteria is noted *S*(*t*)).

Initially the host population is homogeneous, consisting exclusively of sensitive bacteria (except for the data presented in Fig. S2, where all bacteria are initially already CRISPR resistant). Bacteria grow at a maximal rate *r*, but this growth is limited by the density of bacteria (where *k* measures the intensity of density dependence), and bacteria die with mortality rate *m*. At *t* = 0, an inoculum of free Acr-positive phages with density *V*_1_(0) is introduced in the host population, or an inoculum of Acr-negative phages with density *V*_2_(0), or an equal mix of Acr-positive and Acr-negative phages, such that *V*_1_(0) = *V*_2_(0).

All free phage particles adsorb to the bacteria at a rate *a*. When a free phage adsorbs to a sensitive bacterium, two outcomes are possible:

i. with probability 1 − *A*, the phage lyses the host, leading to the release of *B* new phage particles.
ii. with probability *A*, the bacterium acquires CRISPR-based resistance, leading to the destruction of the phage genome.

When an Acr-negative phage particle adsorbs to a CRISPR-resistant bacterium, its genome is destroyed. When an Acr-positive phage adsorbs to a CRISPR-resistant bacterium, three outcomes are possible, as described in our previous model^5^:

i. with probability *ρ*, the Acr-positive phage genome is destroyed prior to expression of the *acr* gene with no change in bacterial resistance (i.e. no immunosuppression). Hence, ρ is a measure of bacterial resistance and increases with the number of spacers targeting the phage. We assume *ρ* is governed by the host and independent of the Acr.
ii. with probability (1 − ρ) *ϕ*, the Acr-positive phage lyses the cell, with the release of *B* new Acr-positive phage particles. Hence, the greater *ϕ*, the greater is the ability of the Acr-positive phage to bypass the CRISPR-Cas immune system.
iii. with probability (1 − ρ) (1 − *ϕ*), the Acr-positive phage fails to complete its lytic cycle but produces some Acr proteins before its genome is cleaved, which blocks bacterial CRISPR-resistance causing the bacterium to become immunosuppressed. This state is reversible and immunosuppressed bacterium become resistant again at rate *γ*. Hence, the smaller *γ*, the longer the bacterium remain in the immunosuppressed state.

If an immunosuppressed bacterium is infected by a phage, the absence of resistance allows the phage to complete its lytic cycle, even if it does not encode an Acr. This yields the following set of ordinary differential equations (see Fig.1A):

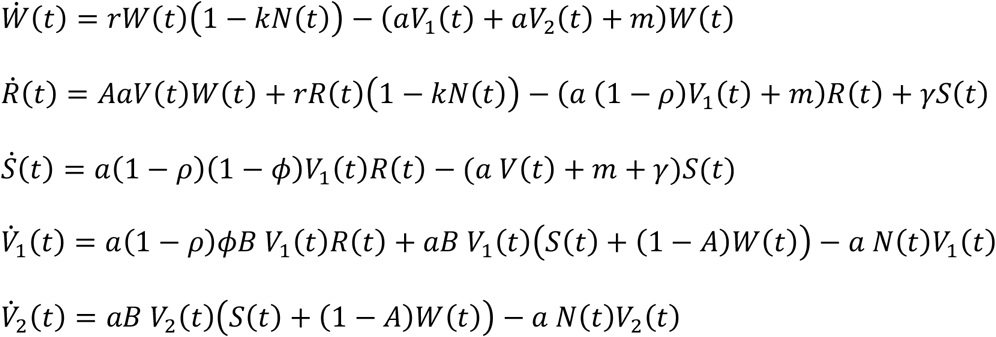

With *N*(*t*) = *W*(*t*) + *R*(*t*) + *S*(*t*), and *V*(*t*) = *V*_1_(*t*) + *V*_2_(*t*).

We used the above model to monitor the infection dynamics following infection with an equal mix of Acr-positive phages (*V*_1_) and Acr-negative phages (*V*_2_) that cannot infect resistant bacteria, with different parameter values *ϕ* and *γ* for the intensity of Acr. All the parameter values are given in the legends of Figs. 1, S1 and S2.

### Bacterial strains and phages

The wild-type strain UCBPP-PA14 of *Pseudomonas aeruginosa* (WT), the derived strain carrying 1 spacer targeting phage DMS3*vir* (BIM-1sp.) and the strain UCBPP-PA14 *csy3::lacZ,* which CRISPR-Cas system is not functional (CRISPR-KO), were used throughout this study. All strains are described in Ref. 5 and references therein. *Escherichia coli* strain DH5α was used to construct and amplify guide-RNA expression plasmids which were subsequently transformed into *P. aeruginosa* PAO1*::spycas9* carrying the *cas9* gene of *Streptococcus pyogenes* under the control of an arabinose-inducible promoter (described in Ref. 23). Strains PAO1*::spycas9*, UCBPP-PA14 WT and CRISPR-KO, but not BIM-1sp., are sensitive to the Mu-like phage DMS3*vir* used in this work (described in Ref. 24). Phage isogenic variants carrying anti-CRISPR genes, namely DMS3*vir*-*acrIF1* and DMS3*vir*-*acrIF4*, were described previously^3,5^. Phage D3112^25,26^, genetically distinct from DMS3*vir* but using the same bacterial receptor (pilus), was used to analyse the evolution of bacterial resistance.

Bacteria were routinely cultured at 37°C either in Lysogeny Broth (LB) broth or M9 minimal medium (22 mM Na_2_HPO_4_; 22 mM KH_2_PO_4_; 8.6 mM NaCl; 20 mM NH_4_Cl; 1 mMMgSO_4_; 0.1 mM CaCl_2_) supplemented with 0.2% glucose. When appropriate (i.e. for plasmids maintenance and expression), LB was supplemented with either 100 µg.ml^−1^ ampicillin (*E. coli* DH5α) or 50 µg.ml^−1^ gentamicin (PAO1*::spyCas9*) and 0.1 % (w/v) arabinose.

### Phage-phage competition experiments

#### Mixed phage infections

Phage mixtures (1:1) of either DMS3*vir-acrIF1* × DMS3*vir* or DMS3*vir* × DMS3*vir-acrIF4* were used to infect fresh cultures of either CRISPR-KO, WT or BIM-1sp. (approx. 4.10^6^ cfu.mL^−1^ in 6 mL of M9 + 0.2% glucose medium, assessed by cell plating), each in 6 replicates. Mixed-infections were carried out at a multiplicity of infection (MOI) of ~0.02 on CRISPR-KO and WT strains and at a MOI of ~50 on BIM-1sp. strain to ensure efficient amplification of DMS3*vir-acrIF1* and DMS3*vir-acrIF4* (DMS3*vir* cannot amplify on BIM-1sp.). Mixed-infections of BIM-1sp. at a MOI of ~0.5 were also performed as negative controls. Cultures were transferred daily (1:100 dilution) into fresh medium and samples were taken at 0, 1, 2 and 3 days post infection (dpi) to monitor the concentrations of each phage population. Upon chloroform extraction (i.e. addition of 1:10 v/v chloroform to cultures, vortex for 30 s and pellet cell debris and chloroform at 4°C), total phage samples were serially diluted in M9 medium and 5 μL of each dilution were spotted on lawns of PAO1::*spycas9* indicator strains (see below for description) that allowed the distinction between competing phages and hence to determine the concentrations of each individual phage population.

#### Generation of PAO1::spycas9 indicator strains to assess concentrations of competing phages

The pJB1 plasmid (described in Ref. 23) harbours a BsaI site for insertion of a desired spacer sequence, a crRNA repeat sequence and a tracrRNA sequence. Upon digestion with BsaI, annealed and phosphorylated oligonucleotides containing the spacer sequence of interest flanked by BsaI sites were ligated into pJB1. PAO1*::spycas9* was transformed with pJB1 expression vectors by electroporation. Briefly, an indicator strain PAO1*::spycas9* carrying a pJB1 expression vector produces a crRNA::tracrRNA-loaded SpyCas9 protein that targets a protospacer complementary to the spacer cloned into the pJB1 vector. As a result, a phage carrying this protospacer cannot produce plaques on that indicator strain. All indicator strains (and corresponding spacer sequences) are listed in table S1.

#### Determination of relative fitness of competing phages by qPCR

Relative frequencies of each phage in the co-culture were measured at 0 and 3 dpi by qPCR, using primer sets specific to each phage, allowing to calculate phages’ relative fitnesses. The following primer pairs were used (designed using Primer Express 3.0.1): ACCTGAGCGAGGATCAATGG and CGCGGCGAACCTTCTG (for DMS3*vir*), CGAAAATGGCAGCAAAATCAA and CCAACGCTTTTGCCGTTT (for DMS3*vir-acrIF1*) and GTGGCGCCCTCCATCAT and GCAAGCCGAAGTAACCATTCTC (for DMS3*vir-acrIF4*). Each pair was tested against the 2 other phages and non-specific amplification could not be detected. Total phage samples obtained upon the above-mentioned chloroform extractions were used as templates. Each qPCR reaction was prepared following manufacturer’s recommendations and composed of 7.5 μl Brilliant III Ultra-Fast SYBR^®^ Green QPCR Master Mix (Agilent Technologies), 0.3 µl of provided reference dye (freshly diluted 1:50 in PCR-grade water, final concentration 300 nM), 1.5 μl primer pair (4.5µM each), 0.15 μl bovine serum albumin (20 mg.ml^−1^) and 3.75 µl undiluted phage sample (or standard phage solution or water), and PCR-grade water to a total volume of 15 µl. For each primer set, standard reactions were performed in triplicate, with six ten-fold dilutions of pure matching-phage stock solutions in PCR-grade water (10^3^ to 10^8^ pfu.ml^−1^), and negative controls (either water or 10^8^ pfus of non-matching phages) were systematically included. Two qPCR reactions were performed on each sample (each reaction being specific to one or the other competing phage). All samples were run at the same time, on a 384-well plate, to avoid between-run measure variations and to ensure that all samples were analysed against the same standards curves. The qPCR program was 95°C for 10 min, followed by 40 cycles of 95°C for 15 s and 60°C for 20 s and was run on QuantStudio™ 7 Flex Real-Time PCR System (Applied Biosystems™). Provided that the efficiencies of the reactions were between 90% and 110%, the threshold cycle (C_T_) was used to calculate the quantity of the targeted-phage in each sample (deduced from standard curves, computed by QuantStudio™ Real-Time PCR Software v1.3). Average quantities (Q) – for the 6 biological replicates of each competition – were then used to calculate phage relative frequencies (fraction, see below), allowing to further calculate phage relative fitnesses (same equation as described in the Bacterial Competition section below).

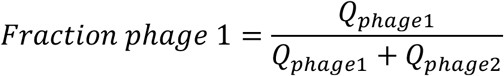

### Individual phage infections

Glass vials containing 3 mL of M9 + 0.2% glucose medium were inoculated with approximately 10^7^ bacterial cells from fresh overnight cultures of WT or BIM-1sp. strains and infected with either phage DMS3*vir*, DMS3*vir*-*acrIF1* or DMS3*vir*-*acrIF4* at initial MOIs of 0.01 (WT) or 100 (BIM-1sp.). Infected cultures were then incubated at 37°C under agitation and transferred daily (1:100 dilution) into fresh medium. Each experiment was performed in 6 replicates. Phage and bacterial concentrations were assessed at T=0, 1 and 3 dpi by performing spot assays and cell plating, respectively.

### Analysis of bacterial evolution of resistance

To determine which phage-immunity mechanisms evolved in WT bacterial populations at 3 dpi, either upon individual or mixed-infections, 24 individual clones (per replicate) were grown in 200 μl of LB broth for 3h at 37°C and 5 μl of these cultures were spotted on LB agar plates, in triplicate. Two microliters (approximately 10^5^ pfus) of either the ancestral phage (i.e. DMS3*vir*, DMS3*vir*-*acrIF1* or DMS3*vir*-*acrIF4*), the alternative phage D3112, or LB broth (negative control) were dropped on top of dried bacterial spots. This phenotypic assay allows to determine whether a given clone is (i) phage-sensitive (lysed by both ancestral and alternative phages), (ii) resistant through surface modification (resistant to both ancestral and alternative phages) or (iii) resistant through CRISPR-immunity (resistant to ancestral phage, sensitive to alternative phage). Presence/absence of CRISPR-Cas-mediated immunity was further confirmed for each clone by PCR using primers CTAAGCCTTGTACGAAGTCTC and CGCCGAAGGCCAGCGCGCCGGTG (for CRISPR array 1) and GCCGTCCAGAAGTCACCACCCG and TCAGCAAGTTACGAGACCTCG (for CRISPR array 2).

### Bacterial competition experiments

Glass vials with 6 mL M9 + 0.2% glucose were inoculated with approximately 2.10^7^ cells from a 1:1 mixture of overnight cultures (grown in M9 medium + 0.2% glucose) either of phage-resistant strains BIM-1sp. and a surface mutant (hereafter referred to as Sm, derived from CRISPR-KO, described in Ref. 7) (Fig. 2D), or of phage-sensitive strains WT and CRISPR-KO (Fig. S3E). Phages (DMS3*vir* or DMS3*vir*-*acrIF1* or DMS3*vir*-*acrIF4*) were then added to each glass vial at a MOI of 0.01 (for the WT × CRISPR-KO competition) or 25 (for the BIM-1sp. × Sm competition). Control competition experiments in the absence of phages were performed in parallel. All bacterial competitions were performed in 6 replicates. Mixed-cultures were transferred daily (1:100 dilution) into fresh medium. At 0, 1 and 3 days after the start of the competition experiment, samples were taken and cells were serially diluted in M9 medium and plated on LB agar supplemented with 50 mg.mL^−1^ X-gal (to allow discrimination between WT or BIM-1sp. (white) and CRISPR-KO-derived (blue) strains). Phage concentrations were also monitored at 0, 1 and 3 days with spot assays. Relative frequencies of competing strains were determined through colony numbers and used to calculate the relative fitness.

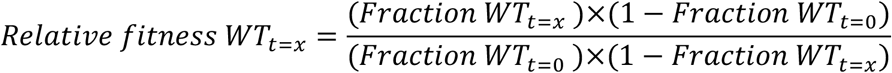

### Amplicon sequencing

Total DNA was extracted from samples of WT bacterial cultures individually infected with either DMS3*vir*, DMS3*vir-acrIF1* or DMS3*vir-acrIF4,* at 3 dpi, using QIAamp DNA Mini kit (Qiagen), according to manufacturer’s instructions. Quality and concentration of DNA samples were verified on a 0.5% agarose gel and assessed by Qubit, respectively. To generate amplicons for subsequent sequencing, the following primers were used: for CRISPR 1 GGCGCTGGAGCCCTTGGGGCTTGG and GCGGCTGCCGGTGGTAGCGGGTG, for CRISPR 2 GCTCGACTACTACAACGTCCGGC and GGGTTTCTGGCGGGAAAAACTCGG. Libraries were prepared by the Centre for Genomic Research (University of Liverpool, UK) and 2×250 bp paired-end reads generated on an Illumina MiSeq platform.

### Bioinformatic analyses

Sequenced reads were trimmed for the presence of Illumina adapter sequences using Cutadapt version 1.2.1^27^. The option -O 3 was used, so the 3′ end of any reads which match the adapter sequence for 3 bp or more are trimmed. Reads were further trimmed using Sickle version 1.2 (https://github.com/najoshi/sickle) with a minimum window quality score of 20. Reads shorter than 20 bp after trimming were removed. Reads were merged with Flash version 1.2.11^28^ and a further 5 bases were trimmed from the 5’ end of each read, following additional quality checks. The resulting read length distributions were determined directly with an Awk expression. Merged reads were then processed using the Qiime2 platform (version 2018.2). Additional quality filtering was done using the default settings based on sequence quality scores (minimum phred score = 4, maximum number of consecutive low scores = 3, minimum length of sequence after filtering = 75% of the original read). Sequences were dereplicated and clustered at 99% similarity using Vsearch^29^. Spacers from the clustered reads were predicted using a modified version of CRISPRDetect^30^ and extracted using a Perl script. Spacers were mapped to the DMS3*vir* genome (based on NCBI_008717 edited to match the sequence described in Ref. 24 using bwa (version 0.7.17) and samtools (1.3.1). The resulting BAM files were plotted in R (version 3.5.1).

### Statistical analyses and graphical output

All graphs of experimental data and statistical analyses were generated with the software GraphPad Prism 7. After verifying that the data (log transformed when appropriate) were not inconsistent with a Gaussian distribution (Shapiro-Wilk normality test), one-tailed t-tests were used to determine whether experimental values (i.e. relative fitness or phage amplification) significantly differed from a theoretical value (e.g. 1), and two-tailed unpaired t-tests were used to compare groups with each other, where appropriate. Each group was composed of 6 values (i.e. 6 independent biological replicates) and 95% confidence levels were used in all statistical tests. In all cases, observed differences were considered significant when p-values were less than a Bonferroni-corrected threshold of 0.05/*a*, where *a* is the number of comparisons.

### Data availability

Amplicon sequencing data have been deposited in the European Nucleotide Archive (ENA) under accession number PRJEB29041. All data generated or analysed during this study are included in this published article (and its supplementary information files).

## Acknowledgments

The authors acknowledge the Centre for Genomic Research (University of Liverpool), in particular R. Eccles, A. Lucaci and R. Gregory for performing amplicon sequencing. The authors thank Dr. L. Zhang for help setting up qPCR analyses and Prof. A. Buckling for critical reading of the manuscript. This work was supported by a grant from the ERC (ERC-STG-2016-714478 – EVOIMMECH), the BBSRC (BB/N017412/1), the Wellcome Trust (109776/Z/15/Z) and the NERC (NE/M018350/1), which were awarded to E.R.W.

## Author contributions

Conceptualization, E.R.W.; Methodology, A.C., S.M., M.L. and E.R.W.; Investigation, A.C., O.F., A.M., SvH; Formal Analysis, A.C., S.M, A.B., S.G; Writing – Original Draft, A.C.; Writing – Review & Editing, A.C., S.M. and E.R.W; Supervision, E.R.W.; Funding Acquisition, E.R.W.

## Materials and correspondence

All materials used in this study are available upon request to Edze Westra or Anne Chevallereau.

## Extended data

**Fig. S1.**
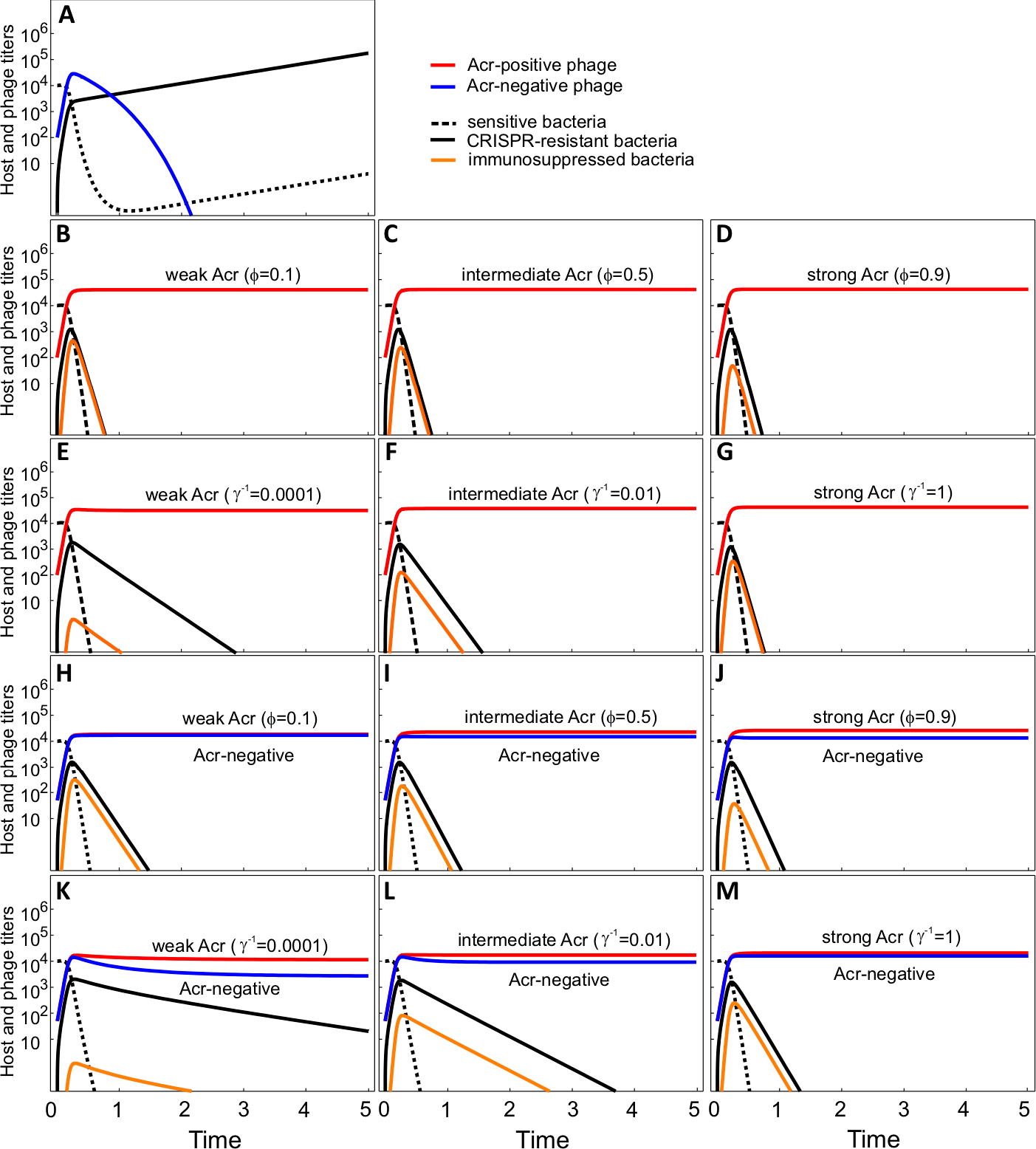
Evolutionary and population dynamics following infection of sensitive bacteria with Acr-negative and Acr-positive phages with different strengths of Acr. (**A**) Infection dynamics and evolution of CRISPR-resistance (solid black line) upon infection of initially sensitive hosts (dashed line) with 100 Acr-negative phages (blue line), and (**B-G**) upon infection with 100 Acr-positive phages (red lines), with different parameter values for *ϕ* and *γ* as follows: (B) *ϕ* = 0.1 and *γ*^−1^ = 1, (C) *ϕ* = 0.5 and *γ*^−1^ = 1, (D) *ϕ* = 0.9 and *γ*^−1^ = 1, (E) *ϕ* = 0.3 and *γ* ^−1^ = 0.0001, (F) *ϕ* = 0.3 and *γ* ^−1^ = 0.01, (G) *ϕ* = 0.3 and *γ* ^−1^ = 1. (**H-M**) Infection dynamics as in (A-G), but upon infection with an equal mix of Acr-negative (blue line) and Acr-positive phage (red line) with different parameter values for for *ϕ* and *γ* as follows: (H)^*ϕ*^ = 0.1 and *γ*^−1^ = 1, (I) *ϕ* = 0.5 and *γ*^−1^ = 1, (J) *ϕ* = 0.9 and *γ*^−1^ = 1, (K) *ϕ* = 0.3 and *γ*^−1^ = 0.0001, (L) *ϕ* = 0.3 and *γ* ^−1^ = 0.01, (M) *ϕ* = 0.3 and *γ* ^−1^ = 1. Other parameter values: a = 0.001, A = 0.2, B = 5, *ρ* = 0.5.

**Fig. S2.**
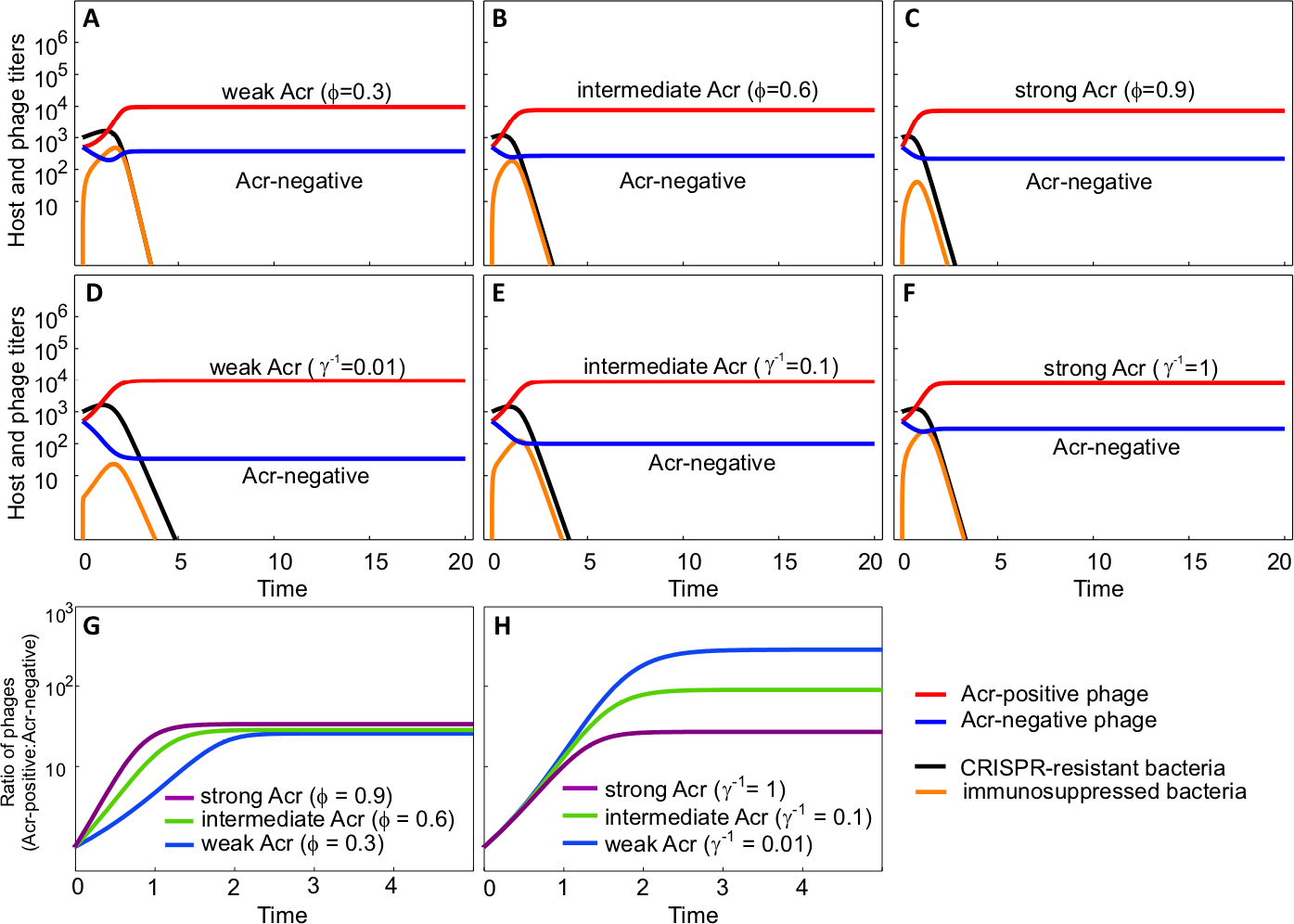
Evolutionary and population dynamics following infection of CRISPR-resistant bacteria with Acr-negative and Acr-positive phages with different Acr strengths. (**A-F**) Population dynamics of CRISPR-resistant bacteria (solid black line), the emergence of immunosuppressed cells (orange line) and phage titres upon infection with an equal mix of Acr-negative phages (blue line), and Acr-positive phages (red line) with different parameter values for *ϕ* and *γ* as follows: (A) *ϕ* = 0.3 and *γ* ^−1^ = 1, (B) *ϕ* = 0.6 and *γ*^−1^ = 1, (C) *ϕ* = 0.9 and *γ*^−1^ = 1, (D) *ϕ* = 0.5 and *γ*^−1^ = 0.01, (E) *ϕ* = 0.5 and *γ*^−1^ = 0.1, (F) *ϕ* = 0.5 and *γ*^−1^ = 1. (**G**) Effect of *ϕ* and (**H**) *γ* on the ratio of Acr-positive and Acr-negative phages following infection of CRISPR-resistant bacteria, with the parameter values of (A-F). Other parameter values: a = 0.001, A = 0.2, B = 5, *ρ* = 0.2.

**Fig. S3.**
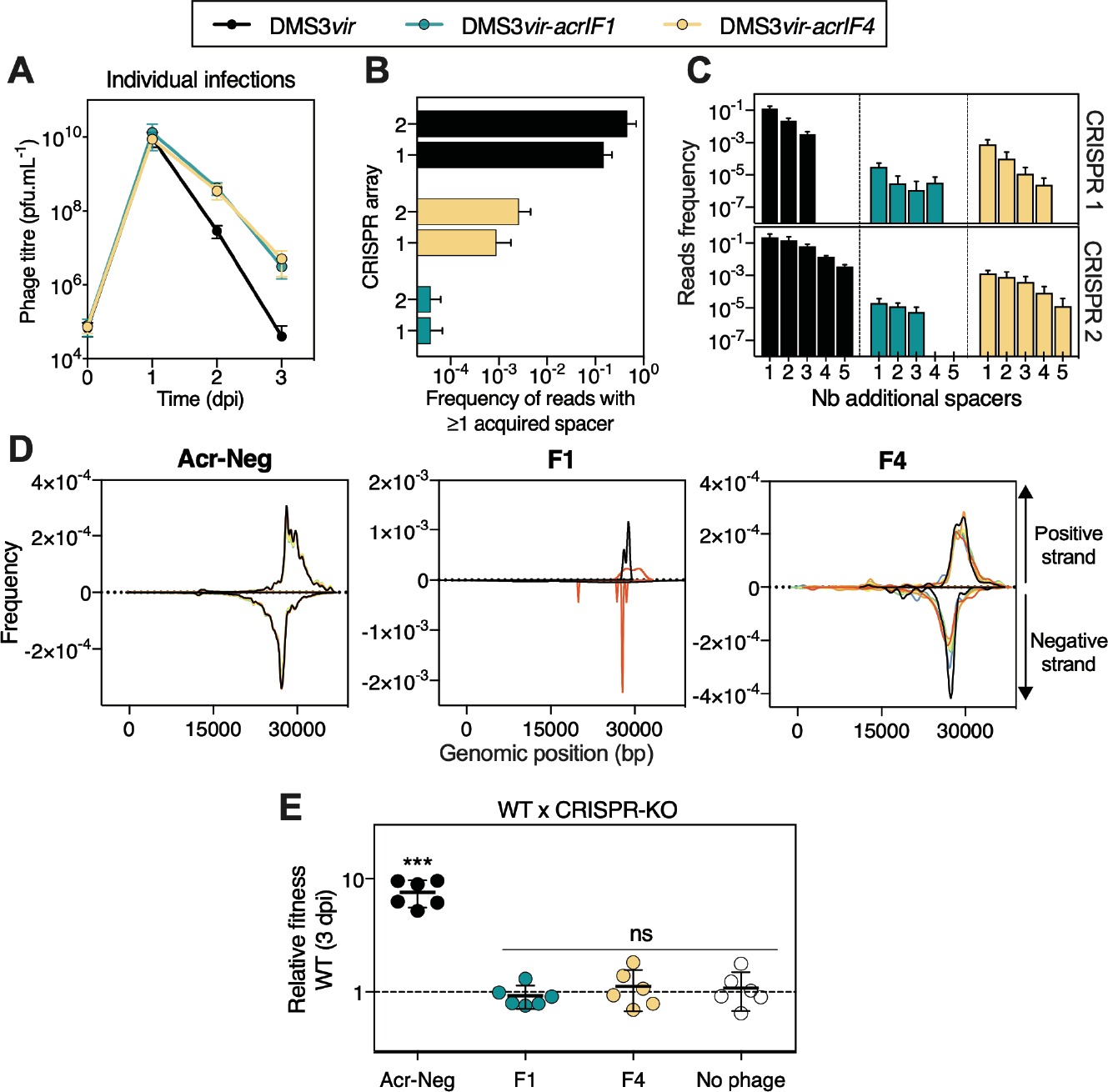
Population dynamics of Acr-positive phages and their impact on the evolution of CRISPR-resistance in the WT host population. (**A**) Phage populations dynamics upon individual infections of the WT host by Acr-negative or Acr-positive phages with different strengths of Acr. (**B**) Frequency of spacer acquisition in CRISPR arrays 1 and 2 at 3 dpi in the evolved WT PA14 populations, as determined by deep sequencing. (**C**) Frequencies of reads containing 1, 2, 3, 4 or 5 additional spacers. (**D**) Protospacer distributions. Newly acquired spacers were extracted from read sequences and corresponding protospacers were mapped back to phage genomes, on positive and negative strands. (**E**) Relative fitness of WT PA14 and CRISPR-KO strains at 3 dpi in the presence or absence of indicated phages at an initial MOI of 0.01. One-sample t-tests indicate significant differences from 1 for ‘Acr-Neg’ (p=0.0004, t_5_=8.3) and no significant differences for ‘F1’ (p=0.42, t_5_=0.87), ‘F4’ (p=0.52, t_5_=0.52) and ‘No phage’ control (p=0.61, t_5_=0.54). In all panels, values for individual replicates and/or means and 95% confidence intervals are indicated. ‘F1’=DMS3*vir*-*acrIF1,* ‘F4’*=*DMS3*vir*-*acrIF4*, ‘Acr-Neg’=DMS3*vir*.

**Table S1.**
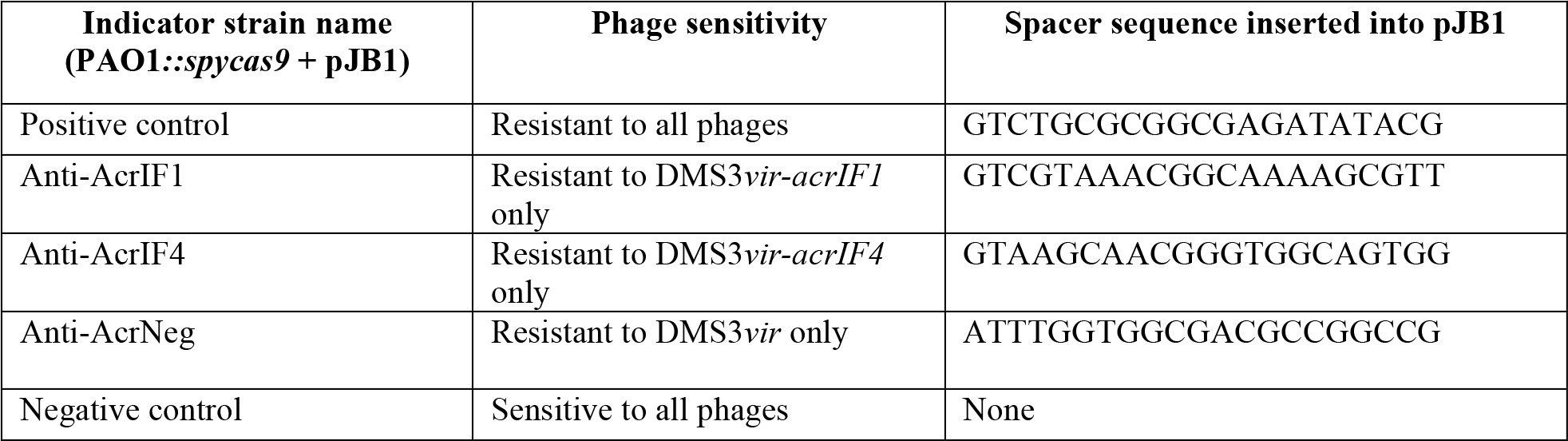
List of PAO1::*spycas9* indicator strains used to monitor individual concentrations of competing phages.

